# Rapid identification of an *Arabidopsis* NLR gene conferring susceptibility to *Sclerotinia sclerotiorum* using time-resolved automated phenotyping

**DOI:** 10.1101/488171

**Authors:** Adelin Barbacci, Olivier Navaud, Malick Mbengue, Rémy Vincent, Marielle Barascud, Aline Lacaze, Sylvain Raffaele

## Abstract

The broad host range necrotrophic fungus *Sclerotinia sclerotiorum* is a devastating pathogen of many oil and vegetable crops. Plant genes conferring complete resistance against *S. sclerotiorum* have not been reported. Instead, plant populations challenged by *S. sclerotiorum* exhibit a continuum of partial resistance designated as quantitative disease resistance (QDR). Because of their complex interplay and their small phenotypic effect, the functional characterization of QDR genes remains limited. How broad host range necrotrophic fungi manipulate plant programmed cell death is for instance largely unknown. Here, we designed a time-resolved automated disease phenotyping pipeline and assessed the kinetics of disease symptoms caused by seven *S. sclerotiorum* isolates on six *A. thaliana* natural accessions with unprecedented resolution. We hypothesized that large effect polymorphisms common to the most resistant *A. thaliana* accessions, but absent from the most susceptible ones, would point towards disease susceptibility genes. This identified highly divergent alleles of the nucleotide-binding site leucine-rich repeat gene *LAZ5* in the resistant accessions Rubenzhnoe and Lip-0. Two LAZ5-deficient mutant lines in the Col-0 genetic background showed enhanced QDR to *S. sclerotiorum*, whereas plants mutated in the closely related *CSA1* gene responded like the wild type. These findings illustrate the value of time-resolved image-based phenotyping for unravelling the genetic bases of complex traits such as QDR. Our results suggest that *S. sclerotiorum* manipulates plant sphingolipid pathways guarded by LAZ5 to trigger programmed cell death and cause disease.

## INTRODUCTION

The fungal pathogen *Sclerotinia sclerotiorum* is the causal agent of *Sclerotinia* stem rot (SSR), also designated as white mold disease, on numerous crop and vegetable species, including rapeseed, soybean, sunflower and tomato. *S. sclerotiorum* can be among the most damaging pathogens of rapeseed and soybean when conditions are favorable (Peltier et al., 2012; Derbyshire and Denton-Giles, 2016). *S. sclerotiorum* penetrates plant tissues through wounds, natural openings, or actively forming compound appressoria (Bolton et al., 2006). It employs a typical necrotrophic strategy to colonize host tissues, rapidly triggering plant cell death (Kabbage et al., 2015; Mbengue et al., 2016). Adapted cultural practices, the use of fungicides and biological control methods are frequently employed to limit damages due to *S. sclerotiorum*, since genetic sources of resistance to SSR are lacking for most crop species (Derbyshire and Denton-Giles, 2016). Instead of a clear demarcation between resistant and susceptible genotypes, plants challenged with *S. sclerotiorum* generally show a continuum of resistance levels designated as quantitative disease resistance (QDR) phenotype (Perchepied et al., 2010; Roux et al., 2014). The molecular bases of QDR in plants remain largely elusive (Poland et al., 2009; Roux et al., 2014). Whereas resistance (R) genes mediating complete disease resistance all belong to the nucleotide-binding site leucine-rich repeat (NLR) family, genes underlying QDR discovered to date span a broad range of molecular functions (Poland et al., 2011; Roux et al., 2014; Corwin and Kliebenstein, 2017). Molecular function of genes associated with QDR include for instance transporters (Krattinger et al., 2009), kinases (Fu et al., 2009; Huard-Chauveau et al., 2013), peptidases (Poland et al., 2011; Badet et al., 2017b) or actin-related proteins (Moscou et al., 2011). NLRs (Debieu et al., 2015; Lee et al., 2016) and signaling components typically associated with R-mediated resistance (Iakovidis et al., 2016) can also mediate QDR, illustrating the tight integration of QDR and R-mediated immunity components.

Because of their complex interplay and their small phenotypic effect, the functional characterization of QDR genes is challenging. With large collections of mutant lines available, studies in the model plant *A. thaliana* enable the rapid functional characterization of individual candidate QDR genes (Huard-Chauveau et al., 2013; Corwin et al., 2016; Rajarammohan et al., 2018). Fully quantitative readouts are often required to reveal the contribution of individual *A. thaliana* genes to QDR against *S. sclerotiorum*, such as ethylene and ROS detection (Perchepied et al., 2010; Zhang et al., 2013), lesion area measurements and estimation of fungal biomass (Zhang et al., 2013; Badet et al., 2017b). To this end, quantitative image analysis can be used as a proxy to evaluate disease severity (Baranowski et al., 2015; Mutka et al., 2016; Badet et al., 2017b; Karisto et al., 2017) and the impact of stress on plant fitness (Chen et al., 2014; Czedik-Eysenberg et al., 2018; Nelson et al., 2018). Increasing the accuracy and robustness of such quantitative phenotyping often involves increasing the number of measurements. This allowed for instance to reveal the role of individual effectors from the bacterial pathogen *Xanthomonas axonopodis* pv. *manihotis* in virulence (Mutka et al., 2016). High-throughput image-based plant phenotyping proved valuable in crop breeding and for research purposes (Araus and Cairns, 2014; Fahlgren et al., 2015; Coppens et al., 2017). There is therefore a need to improve our ability to generate large datasets of quantitative plant disease measurements at low cost, low footprint and reduced human intervention (Czedik-Eysenberg et al., 2018). Because it is generally non-destructive, image-based disease measurement also gives access to the dynamics of disease progression. This enabled for instance to study the spatial and temporal distribution of pathogens in plants tissues (Mutka et al., 2016) and distinguish between infected and non-infected plants long before qualitative symptoms are visible (Czedik-Eysenberg et al., 2018). In spite of remarkable progress in the methods and technology in recent years, fundamental advances in plant pathology enabled by automated plant phenotyping are still relatively limited. For instance, the ability of image-based disease phenotyping methods to identify novel QDR genes and advance our conceptual understanding of this complex trait remains elusive.

Like most pathogens, fungi secrete molecules, often termed “effectors”, to manipulate host physiology and cause disease (Schornack et al., 2009; Doehlemann et al., 2014; Lo Presti et al., 2015). Some pathogen effectors are recognized by specific plant R genes, leading to a rapid and efficient immune response designated as effector-triggered immunity (ETI) (Dodds and Rathjen, 2010). In some plant-pathogen interactions, plant resistance is triggered only in plant genotypes carrying an R-gene enabling the specific recognition of an avirulence effector produced by the pathogen, according to a gene-for-gene model (Flor, 1956). However, this model rarely applies to plant interactions with necrotrophic pathogens. ETI often involves the rapid programmed death of host cells at the site of infection, a process designated as the hypersensitive response (HR) (Mur et al., 2008). Although very efficient to control the spread of biotrophic pathogens, the HR can instead favor the colonization of plants by necrotrophic fungi (Govrin and Levine, 2000). A number of necrotrophic fungal pathogen specialized to infect a few plant species evolved effectors designated as host-specific toxins (HSTs) that trigger cell death specifically in some plant genotypes (Friesen et al., 2008; Oliver and Solomon, 2010). Typical HST examples are the victorin peptides produced by the necrotrophic Victoria Blight fungal pathogen *Cochliobolus victoriae*. Victorin is critical for *C. victoriae* virulence, which is mediated through the specific recognition by the plant LOV1 protein belonging to the NLR class (Lorang et al., 2007). Functional and structural analyses of LOV1 suggest that it corresponds to a typical R gene that has been hijacked by *C. victoriae* to trigger cell death and facilitate infection (Lorang et al., 2012; Wolpert and Lorang, 2016). The wheat *Tsn1* gene belongs to the NLR class and recognizes the ToxA peptide produced by the necrotrophic fungus *Stagonospora nodorum*, conferring effector triggered susceptibility (ETS) to this pathogen (Faris et al., 2010). Another effector produced by *S. nodorum*, SnTox1, triggers programmed cell death when recognized by the wheat gene *Snn1* encoding a wall-associated kinase (WAK) (Liu et al., 2012; Shi et al., 2016). WAKs contribute to resistance against biotrophic and hemibiotrophic fungal pathogens (Hurni et al., 2015; Zuo et al., 2015) providing another example of a biotrophic pathogen defense mechanism hijacked by a specialized necrotrophic fungus. Our knowledge on whether and how broad host range necrotrophic fungi manipulate plant programmed cell death remains nevertheless limited.

The analysis of HR-deficient *Arabidopsis thaliana* mutants suggested that the broad host range necrotrophic fungi *S. sclerotiorum* and *Botrytis cinerea* can benefit from HR cell death (Govrin and Levine, 2000). Similarly, inactivation of the BIK1 kinase enhances *A. thaliana* resistance to avirulent bacterial pathogens but increases susceptibility to *B. cinerea* and *Alternaria brassicicola* (Veronese et al., 2006). Several molecules secreted by broad host range necrotrophic fungi trigger cell death in plants. Examples include *B. cinerea* endo-arabinase BcAra1 (Nafisi et al., 2014), xyloglucanase BcXYG1 (Zhu et al., 2017), the *S. sclerotiorum* necrosis and ethylene inducing peptide SsNEP1 (Dallal Bashi et al., 2010) or *Alternaria tenuissima* Hrip1 protein elicitor (Kulye et al., 2012). Cell death induction by these elicitors is often due to direct toxic effects on plant cells independently of the manipulation of plant programmed cell death (Lenarčič et al., 2017). Conversely, the *A. thaliana* aspartyl protease APCB1 cleaves the BAG6 cochaperone, triggering autophagy and restricting *B. cinerea* colonization (Li et al., 2016). This findings points towards a role for autophagy cell death in resistance to necrotrophic fungi (Lai et al., 2011; Lenz et al., 2011) and highlight the complex role of plant programmed cell death in the interaction with necrotrophic pathogens. Evidence for broad host range necrotrophic fungi exploiting cell death induced by typical plant R genes has not been reported to date.

In this work, we show that the *A. thaliana* NLR gene *LAZ5* (At5g44870), but not its close relative *CSA1*, confers susceptibility to *S. sclerotiorum*. For this, we designed a real-time automated disease phenotyping tool and precisely assessed the kinetics of disease caused by seven *S. sclerotiorum* strains on six *A. thaliana* natural accessions. We found that the speed of lesion growth was highly dependent on the host plant genotype and a good indicator of QDR level. We hypothesized that large effect polymorphisms common to the two most resistant *A. thaliana* accessions, but absent from the four other accessions, would point towards disease susceptibility genes. We identified highly divergent alleles of *LAZ5* in the resistant accessions Rubenzhnoe and Lip-0, while identical alleles existed in the other *A. thaliana* accessions analyzed. As expected, two LAZ5-deficient mutant lines in the Col-0 genetic background showed enhanced QDR to *S. sclerotiorum*, whereas plants mutated in the closely related *CSA1* gene responded like the wild type. Considering that the ectopic activation of LAZ5 triggers cell death (Palma et al., 2010), our results suggest that the broad host range necrotrophic fungus *S. sclerotiorum* exploits this R-gene induced plant cell death to its benefit.

## RESULTS

### Quantification of disease resistance to *S. sclerotiorum* by automated real-time image analysis

To generate massive, quantitative and kinetic measurements of disease caused by *S. sclerotiorum*, we designed mobile imaging cabinets (<u>Nav</u>igable <u>au</u>tomatized phyto<u>tron</u>, Navautron) and the associated automatized image analysis pipeline for detached leaves (**Fig. 1A**). A single Navautron allows imaging simultaneously up to 24 whole *A. thaliana* plants or 120 detached *A. thaliana* leaves. A typical phenotyping session of detached leaves inoculated by *S. sclerotiorum* involves automated imaging every 10 minutes over three days, the automated recognition of disease lesions on individual leaves, yielding a total of >10^6^ lesion measurements with almost no human intervention.

**Figure 1.**
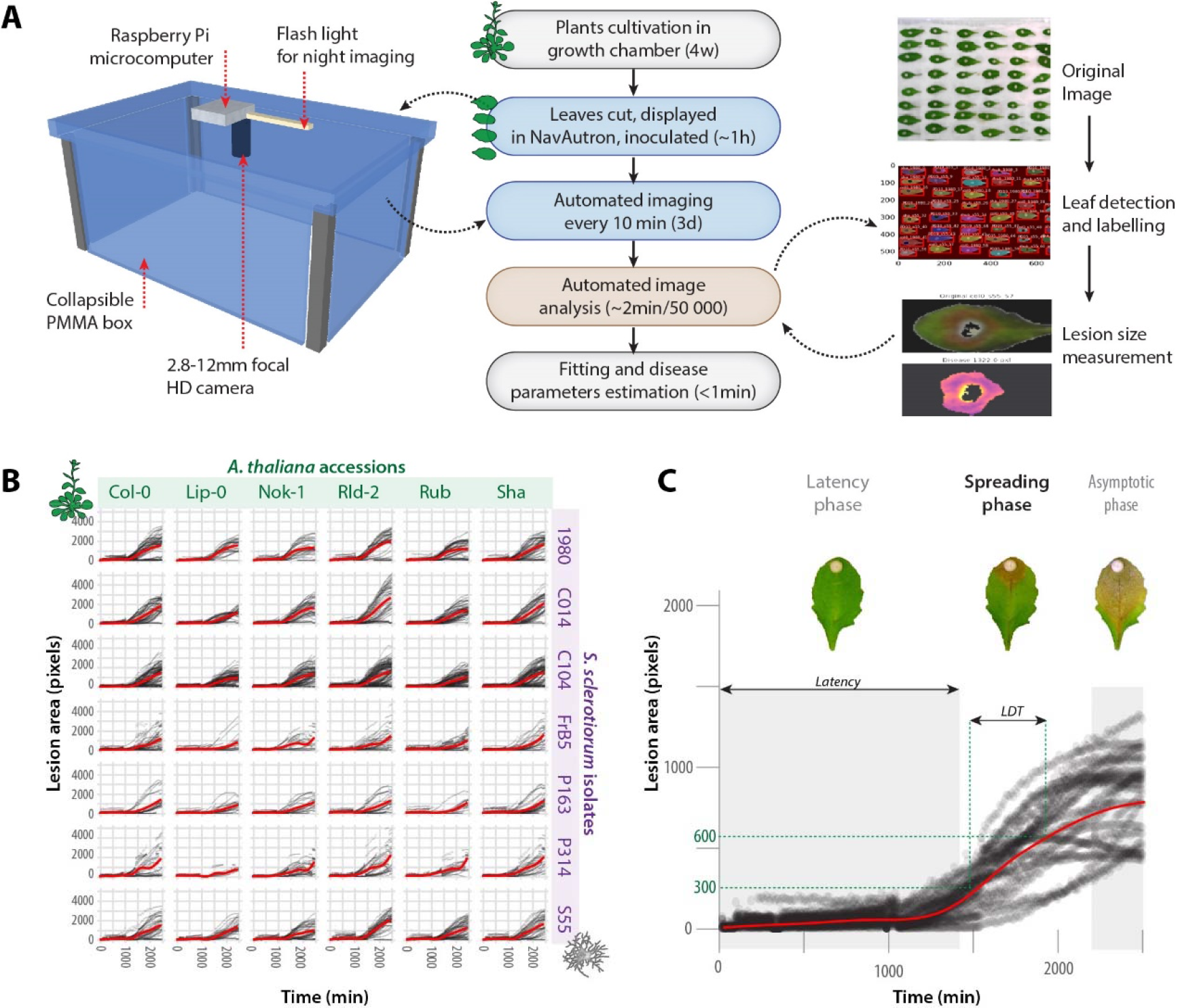
Analysis of quantitative disease resistance (QDR) against ***S. sclerotiorum*** with the Navautron system. **(A)** Overview of the Navautron setup and the pipeline for QDR analysis. Left: Each Navautron consist in a transparent plastic box equipped with a Raspberry Pi microcomputer, a HD camera and a LED flash light. Center: pipeline describing the experiments reported in this manuscript, detached leaves were analyzed with the Navautron through automated imaging, automated image analysis, curve fitting and QDR parameters estimation. The major steps of the automated image analysis are shown on the right. D, days; HD, high definition; PMMA, Poly(methyl methacrylate); w, weeks. **(B)** Kinetics of disease lesion development for 42 different combinations of *A. thaliana* natural accessions (columns) and *S. sclerotiorum* isolates (lines). Red curves show smooth fitting curves for 1,500 to 12,250 measurements. **(C)** Typical kinetics of *S. sclerotiorum* disease lesion development on *A. thaliana*, illustrating the latency phase, spreading phase and asymptotic phase. Characteristic values are the duration of latency phase, dependent on the fungal genotype, and the lesion doubling time (LDT) dependent on the plant genotype.

We quantified the area of necrotic lesions caused by seven isolates of *S. sclerotiorum* (Badet et al., 2017a) on detached leaves from six natural accessions of *A. thaliana* (**Fig. 1B**). We distinguished three major phases along the kinetics of lesions development: (i) a latency phase, during which no necrotic lesion was detected, (ii) a spreading phase, during which the area of necrotic lesions grew exponentially, and (iii) an asymptotic phase, when lesions have reached the border of leaves (**Fig. 1C**). The overall kinetics are therefore depicted by a logistic function. The asymptotic phase was dependent only on leaf size and did not provide information on disease progression. In the 42 interactions tested, the duration of the latency phase and the speed of lesion size increase during the spreading phase were independent (R^2^ = 0.07). The duration of the latency phase was mainly dependent on *S. sclerotiorum* genotype (p=1.2 10^-21^) with a significant but weaker effect of plant genotype (p=2.95 10^-4^). This suggested that the latency phase was primarily determined by fungal strains virulence. Although exponential, the spreading phase can be approximated classically by a linear function through Taylor development at first order. The disease lesion doubling time (LDT, in minutes) is deduced from the slope of the lesion size curve during the spreading phase. The LDT is the characteristic value for the infection since it is sufficient to characterize completely the spreading phase. Of practical importance, the LDT is not dependent on the zoom factor of the cameras used to image symptoms. By contrast with latency, log(LDT) was mostly dependent on the *A. thaliana* genotype (p=7.16.10^-37^) and weakly function of *S. sclerotiorum* isolates (p=2.39.10^-3^). We conclude that the log(LDT) provides an appropriate measure of quantitative disease resistance (QDR) to *S. sclerotiorum*.

### Grouping of plant and fungal genotypes according to QDR phenotypes

Necrotic disease lesions were detected after a latency phase of 1 to 1.5 days in average, mostly determined by *S. sclerotiorum* genotypes. Post hoc pairwise t tests with Benjamini-Hochberg p-value correction (Benjamini and Hochberg, 1995) revealed four groups of *S. sclerotiorum* isolates with distinct latency phases (**Fig. 2A**). Isolates 1980 and p314 had the shorter latency phase (average 22.6 and 23.2 hours), isolates FrB5 and C014 had an intermediate latency phase (average 24.4 and 25.5 hours), isolates C104 and S55 had a longer latency phase (average 26.4 and 26.7 hours) and isolate P163 had the longest latency phase (29.8 hours). The smallest significant difference in latency phase that could be detected in this experiment was about 1h. The ranking of isolates according to their latency phase was identical on all *A. thaliana* accession and therefore not dependent on the plant genotype.

**Figure 2.**
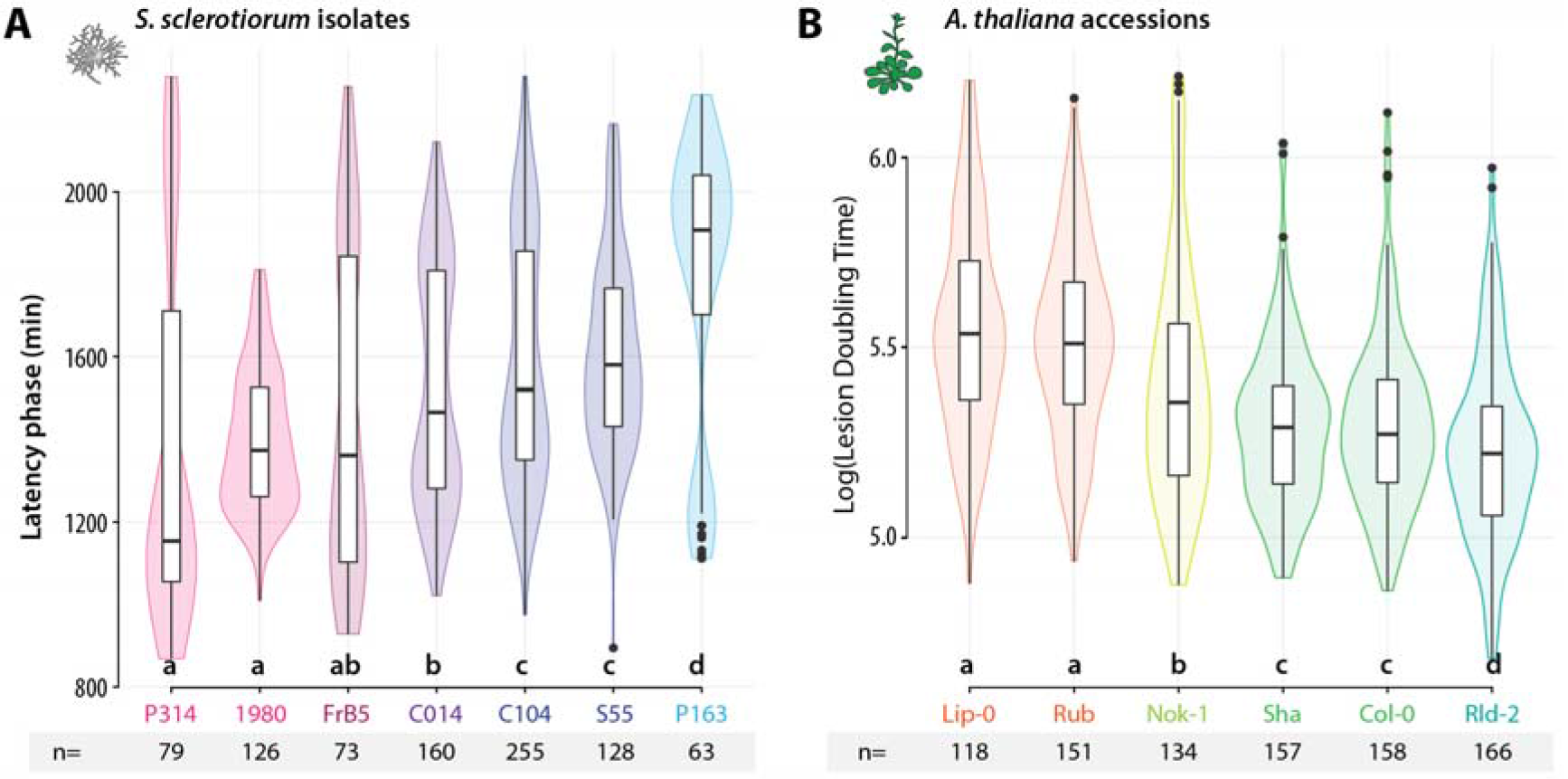
Characteristic values describing disease symptom dynamics in the interaction between seven *S. sclerotiorum* isolates and six *A. thaliana* accessions. **(A)** The duration of latency phase (Y-axis) was mostly dependent on *S. sclerotiorum* isolates (X-axis), ranked from the most (P314) to the least virulent (P163). Duration of the latency phase was measured n=63 to 255 times for each isolate. **(B)** The lesion doubling time (LDT, Y-axis) was mostly dependent on *A. thaliana* accessions (X-axis), ranked from the most (Lip-0) to the least resistant (Rld-2). LDT was measured n=118 to 158 times on each accession. Letters and colors indicate groups of significance determined by post hoc pairwise t tests.

The average LDT ranged between 3.08 and 4.24 hours, corresponding to log(LDT) of 5.22 to 5.54 (**Fig 2.B**), mostly determined by *A. thaliana* genotypes. Post hoc pairwise t tests with Benjamini-Hochberg p-value correction (Benjamini and Hochberg, 1995) revealed three groups of resistance (**Fig. 2B**). With an average Log(LDT) 5.55 and 5.51, Lip-0 and Rubezhnoe (Rub) accessions were substantially more resistant than Nok-1, Col-0 and Shahdara (Sha) (average Log(LDT) 5.40, 5.30 and 5.30 respectively), whereas Rld-2 exhibited the highest susceptibility to any *S. sclerotiorum* strain tested (average Log(LDT) 5.23). The smallest significant difference in LDT that could possibly be detected in this experiment was 13.3 min (Log=0.069). The ranking of *A. thaliana* accessions according to LDT was identical with all *S. sclerotiorum* genotypes. Although the precise ranking of accessions based on LDT differs from the ranking obtained previously with disease severity index on whole plants with *S. sclerotiorum* strains S55 (Perchepied et al., 2010) and 1980 (Badet et al., 2017b), we consistently found Lip-0 and Rub among the most resistant accessions and Rld-2 among the most susceptible ones.

### Identification of candidate susceptibility genes in *A. thaliana* by an association approach

We hypothesized that enhanced QDR in Lip-0 and Rub accessions could result from disruptive mutations in susceptibility genes. To rapidly pinpoint such candidate susceptibility genes, we screened the genome of *A. thaliana* for genes harboring non synonymous mutations both in Lip-0 and Rub accessions but not in any of the other four accessions analyzed (Nok-1, Col-0, Sha and Rld-2) (**Fig. 3A**). For this, we screened 213,624 positions genotyped in the six *A. thaliana* accessions selected (Atwell et al., 2010). We found a total 57,716 single nucleotide polymorphisms (SNPs) present both in Lip-0 and Rub, among which 2,312 were not found in Nok-1, Rld-2 or Sha (unique to Rub and Lip-0). Among those, 298 were non-synonymous SNPs unique to Lip-0 and Rub. The density of non-synonymous SNPs specific to Lip-0 and Rub accessions per gene followed an exponentially decreasing distribution (**Fig. 3B**). Three genes harbored at least four non-synonymous SNPs in Lip-0 and Rub but not any non synonymous SNP in Nok-1, Sha or Rld-2 (**Fig. 3C, Table 1**). This includes *AT1G23935*, encoding an uncharacterized protein with similarity to apoptosis inhibitory proteins, *AT2G33090*, encoding an uncharacterized member of the transcription elongation factor IIS family, and *AT5G44870* encoding *LAZ5*, a disease resistance protein of the TIR-NBS-LRR (NLR) family with similarity to *RPS4* and *CSA1* (Palma et al., 2010). The five non-synonymous SNPs present in the dataset from (Atwell et al., 2010) resided in the NB and the LRR domain of *LAZ5* (**Fig. 4A**). Two additional non-synonymous SNPs are present in the Lip-0 allele of *LAZ5* according to data from http://signal.salk.edu/atg1001/ while the Rub allele harbors large deletions. In a suppressor screen, single point mutations in *LAZ5* resulted in a dominant negative phenotype (Palma et al., 2010), suggesting that Lip-0 and Rub alleles of *LAZ5* could have diverged functionally from their Col-0 counterpart. *LAZ5* is expressed and induced 2.32 fold during the infection of *A. thaliana* by *S. sclerotiorum* (Badet et al., 2017b), and was given the highest priority for functional validation.

**Figure 3.**
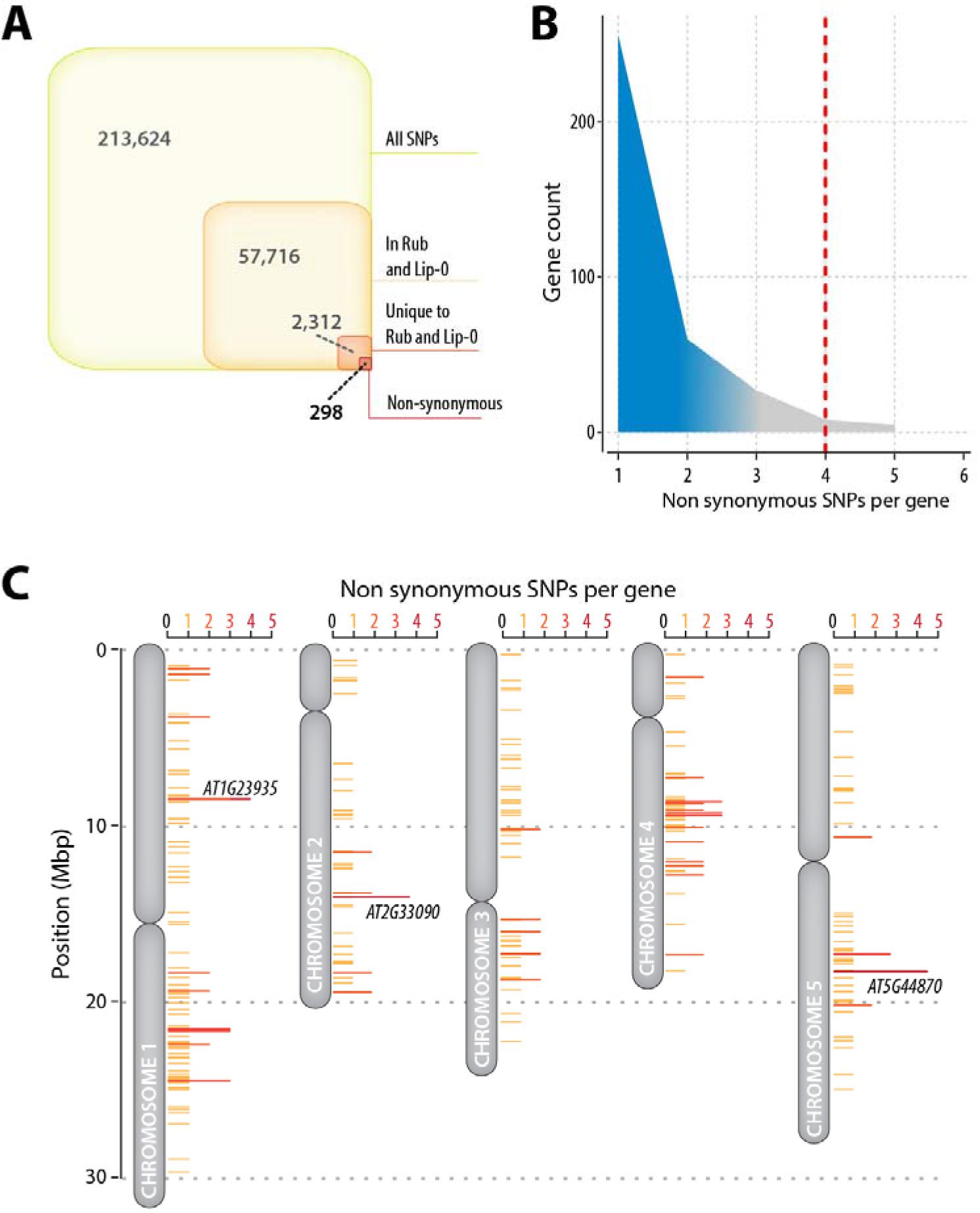
Distribution of single nucleotide polymorphisms (SNPs) in selected *A. thaliana* accessions and identification of candidate disease-relevant genes. **(A)** Distribution of SNPs genotyped by (Atwell et al., 2010) through our pipeline for finding candidate disease-relevant genes. There were 2,312 SNPs common to Lip-0 and Rub but not present in Nok-1, Rld-2 and Sha, among which 298 were non-synonymous SNPs. **(B)** Number of non-synonymous SNP per gene in the list of 298 genes identified in (A). We report on the three genes including at least 4 non-synonymous SNPs in Lip-0 and Rub but no SNP in Nok-1, Rld-2 and Sha (red dotted line). **(C)** Map of A. thaliana chromosomes showing genes with non-synonymous SNPs in Lip-0 and Rub but no SNP in Nok-1, Rld-2 and Sha. Genes discussed in the text are labeled on the figure.

**Figure 4.**
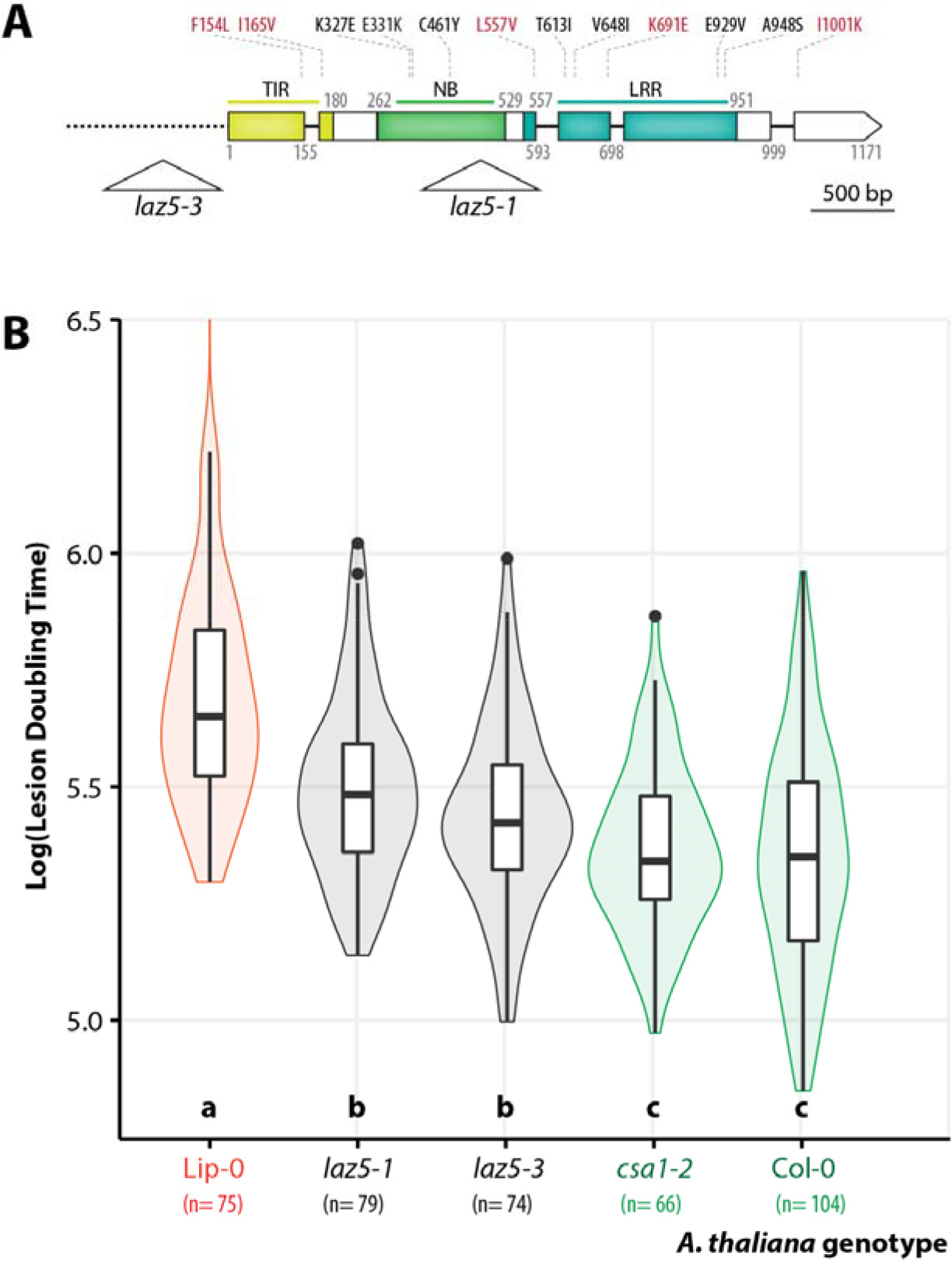
Disruption of the NLR gene LAZ5 reduces lesion doubling time upon *S. sclerotiorum* challenge. **(A)** Schematic map of the *LAZ5* gene showing the position of T-DNA insertion in the *laz5-1* and *laz5-3* mutant lines (triangles). Exons are shown as boxes, introns as plain lines, upstream non coding region as a dotted line. Domains encoded by exons are color-coded and labeled TIR, NB and LRR. Positions are given as amino acid numbers. Non-synonymous mutations known in Lip-0 allele are indicated above boxes, in red when present in (Atwell et al., 2010), in black otherwise. **(B)** Lesion doubling time (LDT, Y-axis) in the most resistant accession Lip-0, two *laz5* mutant lines, the *csa1-2* mutant, and Col-0 wild type. LDT was measured n=74 to 104 times on each accession. Letters and colors indicate groups of significance determined by post hoc pairwise t tests.

**Table 1.** List of genes harboring at last four non synonymous SNPs in Lip-0 and Rub but none in Nok-1, Sha and Rld2. Chr., chromosome; pos. position; LFC, Log_2_ fold gene of gene expression upon inoculation by *S. sclerotiorum*, with the adjusted p-value for differential expression (NA, not applicable).

| Gene id | Description | Chr. | SNP pos. | From | to | SNP type | Gene LFC |
| --- | --- | --- | --- | --- | --- | --- | --- |
| AT1G23935 | Apoptosis inhibitory protein | 1 | 8462159 | C | G | not_syn | 3.93 |
|  |  | 1 | 8462342 | C | G | not_syn | P=0 |
|  |  | 1 | 8463263 | T | C | not_syn |  |
|  |  | 1 | 8465002 | G | A | not_syn |  |
| AT2G33090 | Transcription elongation factor (TFIIS) family protein | 2 | 14035515 | C | G | not_syn | 2.97 |
|  |  | 2 | 14035545 | G | T | not_syn | P=NA |
|  |  | 2 | 14035721 | C | G | not_syn |  |
|  |  | 2 | 14035944 | C | G | not_syn |  |
| AT5G44870 | Disease resistance protein (TIR-NBS-LRR class) family (LAZ5) | 5 | 18115125 | T | C | not_syn | 1.21 |
|  |  | 5 | 18115243 | A | G | not_syn | P=0 |
|  |  | 5 | 18116419 | C | G | not_syn |  |
|  |  | 5 | 18116946 | A | G | not_syn |  |
|  |  | 5 | 18118097 | T | A | not_syn |  |

### Disruption of *LAZ5* increases quantitative resistance to *S. sclerotiorum*

To test for a role of *LAZ5* as a susceptibility gene against *S. sclerotiorum*, we analyzed the phenotype of two *LAZ5* insertion mutant lines in the Col-0 background during infection with *S. sclerotiorum* 1980. The *laz5-1* null mutant (SALK_087262C) carries a t-DNA insertion in the second exon of *LAZ5* and shows dominant suppression of autoimmune cell death in an *acd11-2* background (Palma et al., 2010). The *laz5-3* mutant (SALK_068316) carries a t-DNA insertion ~300bp upstream of *LAZ5* start codon (**Fig. 4A**). Using our Navautron system, we also determined the resistance to *S. sclerotiorum* in *csa1-2* mutant plants defective in *AT5G17880*, a TIR-NBS-LRR gene closely related to *LAZ5* (Faigón-Soverna et al., 2006). As opposed to *laz5-1*, the *csa1-2* mutant showed enhanced susceptibility to avirulent strains of the bacterial pathogen *Pseudomonas syringae* (Faigón-Soverna et al., 2006; Palma et al., 2010). Consistent with our previous measurements (**Fig 2**), the average Log(LDT) was 5.35 on Col-0, similar to the Log(LDT) on the *csa1-2* mutant (Student’s t-test p-value= 0.68) (**Fig 4B**). The *laz5-1* and *laz5-3* mutants had an average Log(LDT) of 5.5 and 5.44, significantly higher than Col-0 (p-value= 1.3e^-05^ and 0.01 respectively). This represented a 16% gain of quantitative disease resistance in the *laz5* mutants compared to wild type. The automated analysis of 74 to 104 plants per genotype thanks to the Navautron setup allowed assessing this quantitative variation robustly. In agreement with our previous measurements (**Fig 2**), the Lip-0 accession showed an average Log_10_(LDT) of 5.7, significantly higher than the *laz5* mutants (p-value = 3.0e^-07^ and 1.6e^-10^). We conclude that disruption of the *LAZ5* gene contributes to the enhanced quantitative disease resistance against *S. sclerotiorum* measured in Lip-0 and Rub accessions compared to Col-0. This identifies *LAZ5* as a TIR-NBS-LRR gene conferring susceptibility to *S. sclerotiorum* through its Col-0 allele.

## Discussion

Quantitative disease resistance is a complex trait governed by multiple genes of small to moderate effect (Poland et al., 2009; Roux et al., 2014; Corwin and Kliebenstein, 2017). Revealing the phenotypic contribution of such small-effect genes challenges our ability to quantify precisely and robustly the level of QDR in diverse plant genotypes. In *Botrytis cinerea* interaction with *A. thaliana*, most plant genes were associated with QDR against a specific *B. cinerea* isolate (Corwin et al., 2016), emphasizing pathogen genetic diversity as a determinant of QDR phenotype. Previous studies used disease index (Perchepied et al., 2010; Rajarammohan et al., 2018), ethylene production (Zhang et al., 2013), camalexin production (Corwin et al., 2016) and lesion area (Corwin et al., 2016; Badet et al., 2017b) to quantify QDR against necrotrophic fungi in natural accessions and mutants of *A. thaliana*. Here we chose lesion area measurement to assess QDR for being (i) a continuous parameter (by contrast to discrete disease indices), (ii) non-destructive and therefore giving access to disease kinetics, and (iii) amenable to automated image-based measurement, opening the way to user-independent high-throughput quantification. For the experiments reported in this manuscript, we conducted *S. sclerotiorum* inoculations and QDR measurement on *A. thaliana* detached leaves to reach 120 samples analyzed with a single Navautron cabinet. This approach does not provide a complete characterization of plant QDR as discrepancies may exist between detached-leaves and whole plant disease resistance (Liu et al., 2007), lesion area may differ from area colonized by the fungus (Kabbage et al., 2015). This could explain differences in the ranking of *A. thaliana* accessions according to resistance between this study and (Perchepied et al., 2010). Our Navautron pipeline gives access to the kinetics of symptoms development during plant-fungal pathogen interactions, which offers several advantages over single end-point measurements. First, it allows uncoupling lesion doubling time (LDT) and latency duration. Since latency appeared mostly independent from plant genotype in our analysis, LDT provides a more direct measurement of plant QDR potential than end-point measurements. This approach would allow untangling plant and pathogen genetic factors contributing to quantitative immunity (Corwin et al., 2016). Second, the resolution and data richness offered by real time image-based phenotyping allows detecting small phenotypic effects that are not accessible to classical approaches, revealing for instance the virulence function of single pathogen effectors (Mutka et al., 2016). Here we could detect robustly variations as small as 1h (about 3.9%) in latency period and 12 minutes (about 5.24%) in LDT. Finally, LDT is independent of leaf shape and size and allows comparison of QDR in plants with contrasted leaf architectures. In this initial study, *A. thaliana* accessions ranked identically for their LDT against seven *S. sclerotiorum* isolates, arguing against specific plant-pathogen genotype interactions below the species level. Previous studies reported genetic determinants of plant QDR specific of pathogen genotypes (i.e evidence for “pathotypes”) in the interaction of *B. cinerea* with *A. thaliana* interaction (Corwin et al., 2016) and *S. sclerotiorum* with *Brassica napus* and *B. juncea* (Ge et al., 2012; Barbetti et al., 2014). These pathotypes may result from the combined effect of fungal genetic determinants of latency duration and plant determinants of LDT. Future experiments will expand the diversity of *S. sclerotiorum* isolates and *A. thaliana* accessions screened by real time phenotyping to search for specific plant x pathogen interactions affecting LDT.

Time-resolved image-based phenotyping with the Navautron system allowed identifying *LAZ5* as a susceptibility gene to *S. sclerotiorum. LAZ5* belongs to the TIR-NBS-LRR family and is required for ACD11-mediated cell death (Palma et al., 2010). Overexpression of *LAZ5* results in hypersensitive cell death, while *LAZ5* mutant alleles suppress the autoimmune phenotype of *acd11* mutants (Palma et al., 2010). To our knowledge, this analysis provides with *LAZ5* the first example of a TIR-NBS-LRR gene controlling susceptibility to a broad host range necrotrophic fungus. Whether effector molecules secreted by *S. sclerotiorum* interfere directly or indirectly with *LAZ5* function remains to be determined. Testing for enhanced resistance of *laz5* mutants to diverse *S. sclerotiorum* isolates and necrotrophic fungal species should be a promising future direction to address this question.

Broad host range fungal pathogen invest a substantial fraction of their cellular energy in secreted proteins to counter plant defenses and extract nutrients from plant cells (Badet et al., 2017a). Manipulation of the *LAZ5* pathway may be part of *S. sclerotiorum* strategy to actively trigger plant cell death. Autoimmune mutants such as *acd11* were proposed to be altered in functions guarded by NB-LRR genes such as *LAZ5* (Palma et al., 2010; Tong et al., 2017). *ACD11* encodes a ceramide-1-phosphate transfer protein, which led to the hypothesis that LAZ5 could guard sphingolipid metabolic pathways targeted by pathogen effectors (Simanshu et al., 2014). Necrosis and ethylene-inducing peptide 1–like (NLP) are toxins secreted by diverse plant pathogens that target plant sphingolipids (Lenarčič et al., 2017). *S. sclerotiorum* produces NLPs (Dallal Bashi et al., 2010) the activity of which could be guarded by *LAZ5. A. thaliana* mutants in the dihydrosphingosine-1-phosphate lyase AtDLP1 are more resistant to *B. cinerea* (Magnin-Robert et al., 2015) supporting the view that sphingolipids are important mediators of QDR against necrotrophic fungi. Alternatively, LAZ5 activity could be costly for plants so that the inactivation of this gene would be beneficial even in the absence of pathogen. However, we did not detect growth defects in *laz5* mutants, arguing against this hypothesis. The similar LDT measured in *csa1-2* and wild type plants upon *S. sclerotiorum* inoculation also pleads for a relatively specific role of *LAZ5* in QDR against this fungus. *CSA1* and *RPS4* are two TIR-NBS-LRR genes closely related to *LAZ5* that confer resistance to avirulent strains of the bacterial pathogen *Pseudomonas syringae* (Gassmann et al., 1999; Faigón-Soverna et al., 2006), whereas *LAZ5* does not (Palma et al., 2010), supporting LAZ5 specific function.

Inactivation of *LAZ5* contributed ~30% of the reduced LDT in Lip-0 accession, indicating that Lip-0 harbors other genetic variants positively affecting QDR. Lip-0 also shows high QDR against *B. cinerea, Plectosphaerella cucumerina* and *Fusarium oxysporum* (Llorente et al., 2005), making it a useful natural resource for studying the determinants of QDR. The Navautron system is flexible allowing for variations and improvements. Future experiments will include imaging whole plants to detect putative spatial bottlenecks to disease progression, age-related factors associated with QDR and tradeoffs between QDR and plant growth.

## MATERIAL AND METHODS

### Plant material and cultivation

Six natural accessions of *A. thaliana* were chosen to cover the range of resistance to *S. sclerotiorum* (Perchepied et al., 2010): Lip-0 (CS76542; ecotype ID: 8325), Rubenzhnoe-1 (CS76594; 7323), Nok-1 (CS78282; 7270), Shahdara (CS78397; 6962), Col-0 (CS76778; 6909), Rld2 (CS78349; 7457). Plants were grown at 22°C, 9 hours light period at 120 μmol/m2/s during 4 weeks before inoculation. Arabidopsis insertion mutant lines in the Col-0 background were obtained from the Nottingham Arabidopsis Stock Centre (NASC). Mutant lines in *LAZ5* (*AT5G44870*) were SALK_087262C (*laz5-1*) and SALK_068316 (*laz5-3*), the csa1-2 mutant line was SALK_057697C. Homozygous T-DNA insertion was verified by PCR for each line following recommendations from the NASC website.

### Fungal cultivation and inoculation

The seven isolates of *S. sclerotiorum* used in this work are described in (Badet et al., 2017a). Four isolates (C014, C104, P314, P163) were obtained from a rapeseed field population collected in Blois (France) in 2010, isolate FrB5 was obtained from (Vleugels et al., 2013), isolate S55 was obtained from (Perchepied et al., 2010) and isolate 1980 is S. sclerotiorum reference strain (Derbyshire et al., 2017). *S. sclerotiorum* isolates were grown on Potato Dextrose Agar (PDA) plates for 5 days at 23°C in the dark prior inoculation. 240 combinations of *A thaliana* accessions and *S sclerotiorum* isolates were distributed in two Navautrons in a fully randomized design. Leafs were cut with a scalpel blade at the time of inoculation, placed adaxial face up on wet paper towel overlaid at the bottom of Navautrons. Inoculations were performed as described in (Badet et al., 2017b), using PDA plugs of 5mm diameter colonized by the fungus placed upside down on the adaxial surface of leaves (mycelium in contact with the leaf). The experiment was repeated three times independently and results from the three replicates were combined for analysis. Statistical analyses were performed using R software and the ‘car’ library for type II ANOVA and pairwise t test with Benjamini and Hochberg p-value correction (Benjamini and Hochberg, 1995; R Core Team, 2014). Distributions of LDT were normalized by log transformation. Plots were made using the ‘ggplot2’ library (Wickham, 2010).

### Design of the Navautron phenotyping cabinets

Five mm-wide PMMA plates were cut on a Trotec Speedy 500 Laser cutting machine according to plans provided as **Supplementary file 1**. Holes were drilled at the center of the upper panel to place full HD 1080p USB cameras with 2.8-12mm focal length (model ELP-USBFHD05MT-FV-F1 manufactured by ELP, China). The cameras were plugged to Raspberry Pi 3 Model B motherboards equipped with Waveshare 3.2 inch TFT touchscreens. Autofocus and automatic white balance were disabled. For each experiment, pictures were taken every 10 minutes during 4 days and stored on SD cards. During nighttime, a LED flash light is turned on during 5 seconds for image acquisition. Navautrons were placed in Percival AR-41L2 growth chambers at 22°C with ~90% humidity under 9h light period.

### Image analysis pipeline

The area of necrotic disease lesions was computed by numeric method for each individual leaf at each time point. We designed is a thresholding method based on color analysis using the scikit-image python image analysis toolbox (van der Walt et al., 2014). The method splits images of an infected leaf into three layers corresponding to the background, the lesion and the leaf. The ‘background’ corresponded to pixels with a saturation value superior or equal to 0.3 in HSV color space. The lesion corresponded to pixels with red component higher than green value in RGB color space. The area of disease lesions was computed as the sum of pixels in the ‘lesion’ layer. The remaining pixels were attributed to the ‘leaf’ layer. Image analysis allowed the collection of lesion area values over time. Every kinetics was fitted by a polynomial regression. The latency time and lesion doubling time were extracted from the fit. The latency time was defined as the time needed to reach a lesion area of 300 pixels. The lesion doubling time was computed as the difference between the time when lesion area reached 600 pixels and the time when lesion area reached 300 pixels. Kinetics that exhibited excessive signal to noise ratio were automatically excluded from the analysis as follows: we excluded leaves of size <300 pixels and leaves with a maximum lesion size < the variance of lesion size during the latency period. About 3 leaves were excluded per Navautron representing <2.5% of all leaves.

### Analysis of polymorphisms and identification of candidate genes

We used a dataset of 214,000 SNPs on 1,196 *A. thaliana* accessions (Atwell et al., 2010; Horton et al., 2012) to search for non-synonymous mutations present in Lip-0 and Rubezhnoe accessions but not in Col-0, Nok-1, Shahdara and Rld-2. The functional consequences of SNPs were predicted automatically using a script derived from the BioPython library (Cock et al., 2009) provided as **Supplementary file 2**. For this, gene models from TAIR version 9 were used. Predictions were verified manually for the top candidates using ExPASy translate tool and http://signal.salk.edu/atg1001/ genome browser.

## Supporting information

## Authors’ Contributions

A.B., O.N. and S.R. conceived the original screening and research plans; A.B., O.N., M.M and S.R. supervised the experiments; R.V., M.B and A.L. performed most of the experiments; A.B. and S.R. designed the experiments and analyzed the data; A.B and S.R. conceived the project and wrote the article with contributions of all the authors; A.B and S.R. agree to serve as the authors responsible for contact and ensure communication.

## Accession Numbers

Arabidopsis Genome Initiative locus identifiers for the genes in this article are as follows: At5g44870 *(LAZ5);* AT5G17880 *(CSA1)*.

## Acknowledgements

We are grateful to past and present members of the QIP lab for stimulating discussions and suggestions. This work was supported by a starting grant of the European Research Council (ERC-StG 336808 project VariWhim) to S.R. and the French Laboratory of Excellence project TULIP (ANR-10-LABX-41; ANR-11-IDEX-0002-02).

