## Supplementary figures and images for "Rapid identification of an *Arabidopsis* NLR gene conferring susceptibility to *Sclerotinia sclerotiorum* using time-resolved automated phenotyping"

### Supplementary file 1

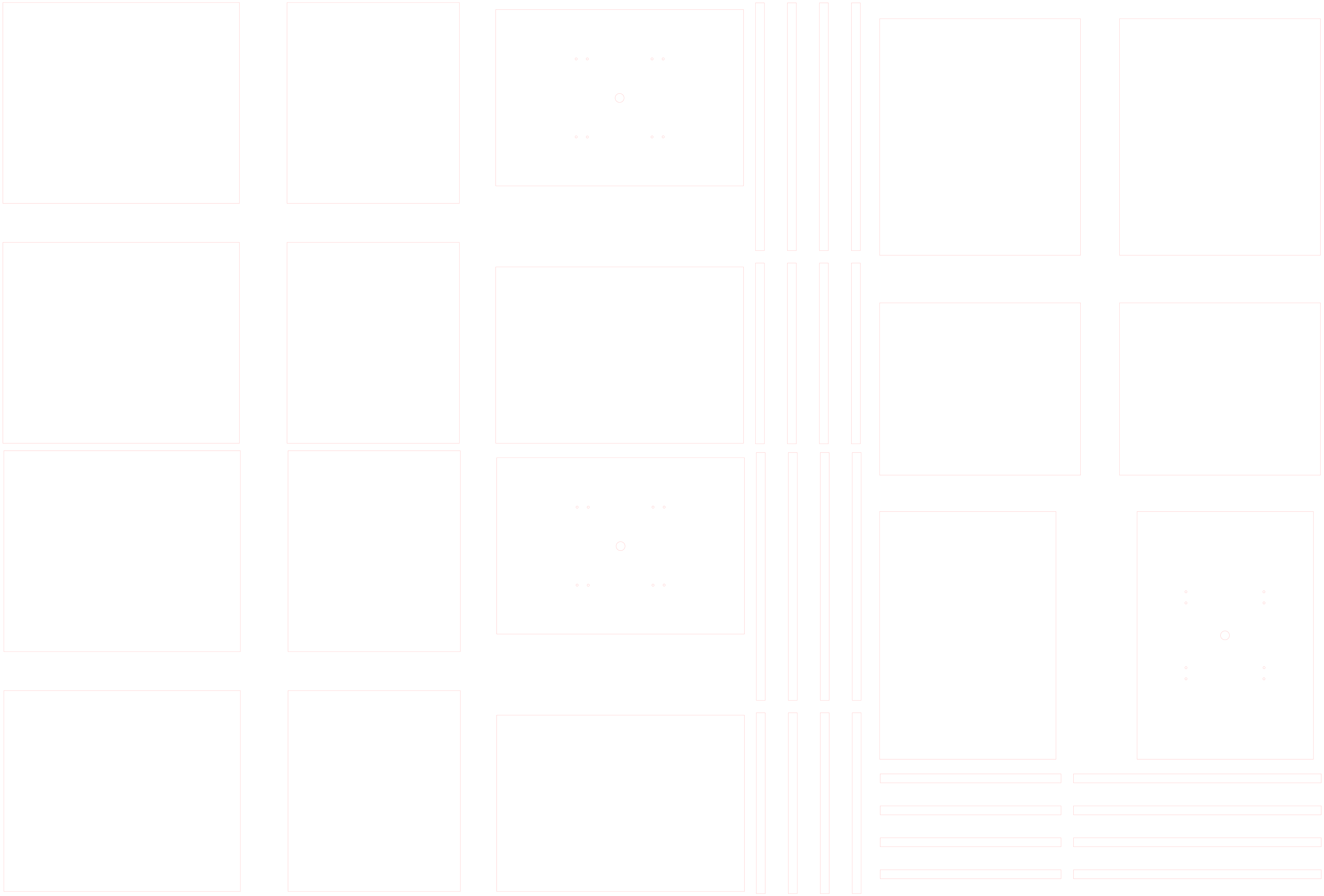

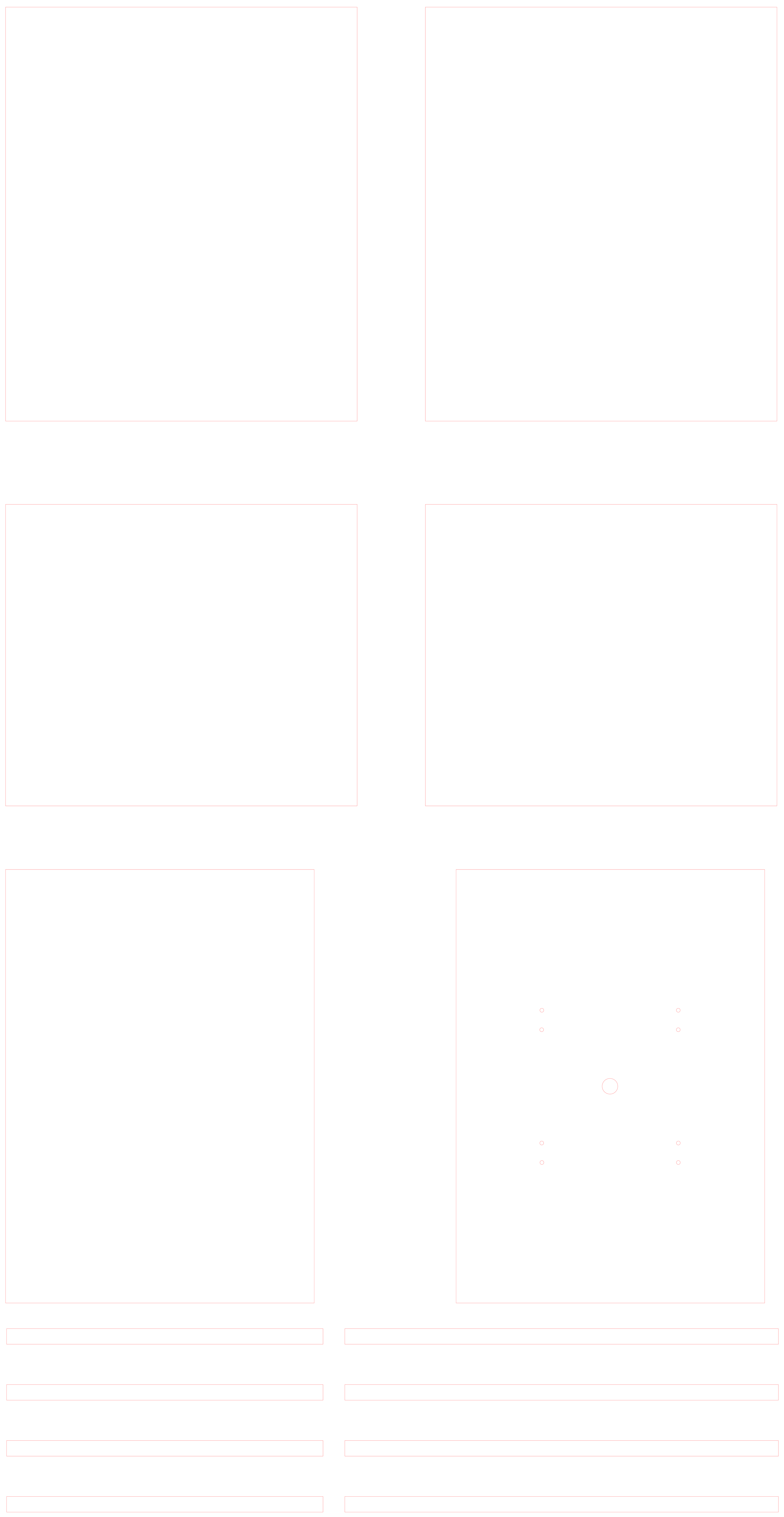
